# Negative emotional visual stimuli alter specific improvised dance biomechanics in professional dancers

**DOI:** 10.64898/2026.03.18.711707

**Authors:** Bruno Cesar Burin Maracia, Thales Rezende Souza, Gabriela Silva Oliveira, Júlia Beatriz Palma Nunes, Carolina Eduarda Santiago dos Santos, Camila Batista Peixoto, Júlia Beatriz Lopes Silva, Líria Akie Okai de Albuquerque Nóbrega, Priscila Albuquerque de Araújo, Renan Pedra Souza, Bruno Rezende Souza

## Abstract

Dance is a core form of human-environment interaction and a powerful medium for emotional expression, yet dancers are routinely exposed to environmental affective cues that may shape their movement. We tested whether a negative emotional context induced immediately before improvisation alters dance biomechanics. Twenty professional dancers performed two 3-min improvised dances. Between dances, they viewed either Neutral or Negatively valenced pictures from the International Affective Picture System (IAPS; 2 min 40 s, 5 s per image). Eye tracking verified attention to the visual stream. Mood was assessed at four time points (PT1–PT4) using the Brazilian Mood Scale (BRAMS), and full-body, three-dimensional kinematics were captured at 300 Hz using a 9-camera optoelectronic system (Qualisys) and processed to measure global movement amplitude and expansion. Negative IAPS exposure increased tension, depression, fatigue, and decreased vigor from PT2 to PT3. Biomechanically, the Negative Stimulus dancers showed a significant reduction in global movement amplitude after negative IAPS exposure, with reduced movement amplitude of the body extremities. In contrast, global movement expansion remained unchanged; that is, the extremities were not positioned closer or farther from the pelvis. Neutral images produced no mood change and no measurable modulation of movement amplitude or expansion. Together, these results support the hypothesis that improvised dance carries biomechanical signatures of the dancer’s current affective state, beyond the intended expressive content, and provide an automated motion-capture workflow for studying emotion-movement coupling in spontaneous dance.

**Highlights:** Negative visual context shifted dancers’ mood toward negative affect

Negative images reduced movement amplitude in improvised dance

Movement expansion remained stable despite mood induction

**Graphical Abstract:** 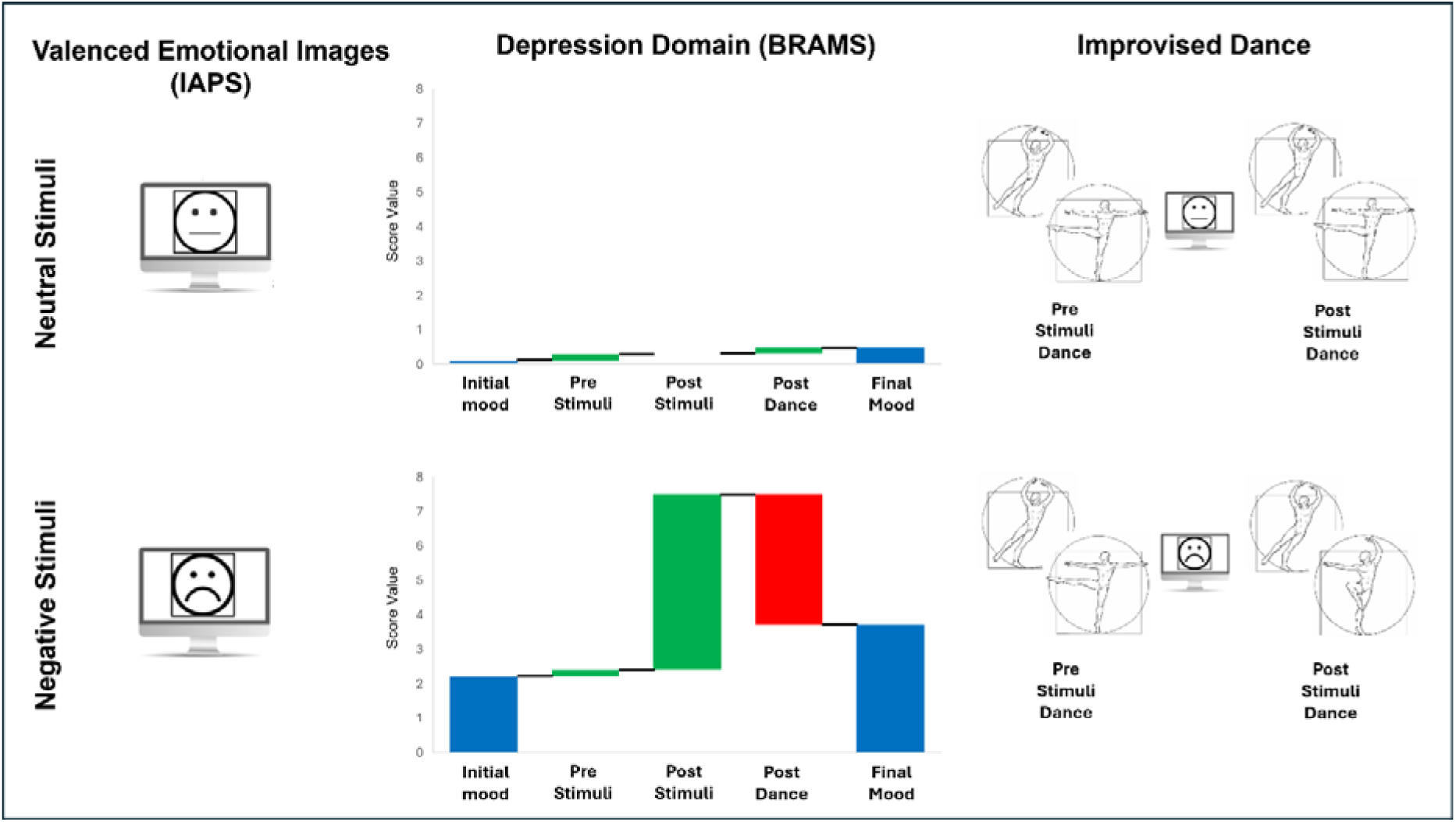

---

Artists routinely externalize internal states through art, translating fleeting emotions into sounds, images, words, and movement. Yet a central question remains: to what extent does an artwork reflect the emotion an artist intends to convey rather than the emotion the artist is currently experiencing? Few studies across creative domains suggest that affective state can bias multiple stages of artistic production, from idea generation to expressive execution. For example, affect (especially positively valenced, activating states) can measurably shift creative performance, underscoring that creativity is not affect-neutral (Baas et al., 2008; He, 2023; To et al., 2012; Verhaeghen et al., 2005). In music, performers can also generate improvisations in response to emotional cues, with quantifiable changes in structural features, supporting the idea that affective context can be “written into” the product itself (McPherson et al., 2014).

This affect-behavior coupling is not limited to artistic outputs. It generalizes movement-based activities in which the quality and efficiency of action matter. In sport and other high-skill contexts, negative affect and anxiety can impair performance, often by altering attention, control strategies, and the balance between automatic and consciously monitored execution (Horikawa & Yagi, 2012; Nibbeling et al., 2012; Wilson, 2008; Yu, 2015). More broadly, affective stimuli can produce rapid, measurable modulations in motor output. Laboratory studies using emotionally evocative pictures show speed-accuracy trade-offs during motor tasks, suggesting that emotional processing can directly influence movement attributes (Coombes et al., 2005). Complementary work demonstrates that emotional context shapes postural control and gait initiation kinematics, consistent with the notion that affect primes action tendencies and alters motor preparation (Benesova et al., 2026; Stins & Beek, 2007). Even goal-directed reach-to-grasp actions can show valence-dependent tuning of temporal kinematic features, indicating that emotional meaning can propagate into the biomechanics of everyday movements (Esteves et al., 2016).

Dance sits at the simultaneous intersection between a creative practice and a complex motor behavior. Improvised dance, in particular, offers a naturalistic window into how affect may influence movement without the constraints of pre-learned choreography. Recent motion-capture studies have quantified kinematic correlates in unconstrained improvised dance, establishing that rich biomechanical structure can be extracted from spontaneous movement (Hartmann et al., 2019). Moreover, spontaneous dance has been linked to post-dance changes in experienced affective valence, reinforcing the bidirectional relationship between emotion and improvised movement (Bernardi et al., 2017). Together, these findings motivate a mechanistic question: Does negative emotional context systematically reshape the biomechanics of improvised dance in ways detectable beyond subjective report?

An approach to experimentally controlling affective context is the International Affective Picture System (IAPS), a standardized set of images with normative affective ratings that has become an important tool of affective science (Lang et al., 2020). IAPS stimuli reliably elicit changes in subjective valence and arousal, as well as in psychophysiological responses. It has been widely adopted and culturally adapted (including norms in Brazilian samples and European Portuguese adaptations), enabling reproducible emotional manipulations across laboratories and populations (Branco et al., 2023; R. L. Ribeiro et al., 2004). Importantly, IAPS is not only suitable for probing perception and autonomic responses, but it is also well-suited for testing motor consequences of emotional stimulation, because exposure to affective pictures has been repeatedly shown to alter postural control, gait initiation, and other movement parameters (Bouman et al., 2015; Stins & Beek, 2007).

Building on evidence that affective states can modulate creative performance (Baas et al., 2008; He, 2023; To et al., 2012; Verhaeghen et al., 2005), shape the structure of improvisational products (McPherson et al., 2014), and modulate motor behavior (Coombes et al., 2005), we hypothesize that improvised dance movements reflect not only the emotion a dancer intends to communicate, but also the dancer’s momentary affective state. Accordingly, beyond choreographic or expressive intent, the biomechanics of improvisation may carry an additional emotional “signature” imposed by transient emotional context. To test this, we use normatively rated images from the IAPS to induce negative affect and quantify its effects on spontaneous dance biomechanics (Esteves et al., 2016; Hartmann et al., 2019).

## Method

### Participants

Professional dancers were recruited via social media, email, and WhatsApp outreach. The final cohort comprised 20 participants (40% male, 60% female), randomly assigned to either the Neutral Stimuli group (n = 10), exposed to neutral visual stimuli, or the Negative Stimuli group (n = 10), exposed to negative emotional visual stimuli. The study employed a double-anonymized design, with neither the participants nor the principal investigator aware of group assignments. Inclusion criteria for participation in the study were: (1) over 18 years of age; (2) professional salaried dancer with at least 5 years of experience; (3) active as a dancer in group or individual performances; (4) no severe ocular pathologies; (5) availability for in-person data collection. Exclusion criteria included: (1) inability to perform the dance task; (2) inability to comprehend and execute the general research instructions.

### Study Design and Procedures

This randomized controlled study was conducted in a temperature-controlled environment (21-23°C). Participants first completed a demographic and eligibility questionnaire, which included an assessment of limb dominance. Subsequently, 58 passive reflective markers were affixed to anatomical landmarks in accordance with the van Sint Jan protocol (Jan, 2007). Movement was tracked using a 9-camera, 3D motion capture system (Oqus 7 cameras, Qualisys) with two marker clusters placed on the thigh and calf of each leg.

The experimental procedure began with an initial assessment of participants’ emotional state using the Brazilian Mood Scale (BRAMS), a 24-item questionnaire that evaluates six mood domains (Psychometric Test 1: PT1) (Bevilacqua et al., 2019; Searight & Montone, 2017; Shacham, 1983; Terry et al., 1999). Following a brief habituation phase, participants performed a 3-minute dance improvisation using everyday movements, under standardized instructions provided by the researcher (Issartel et al., 2017; M. M. Ribeiro, 2017). Immediately after the improvisation, they completed the BRAMS for a second time (Psychometric Test 2: PT2). Participants were then randomly assigned to either the Neutral Stimuli group or the Negative Stimuli group. The Neutral Stimuli dancers were exposed to neutral visual stimuli, while the Negative Stimuli dancers were exposed to negative affective visual stimuli. The stimuli consisted of 32 images from the International Affective Picture System (IAPS), each displayed for 5 seconds on a 36 x 96 cm monitor positioned 1 meter from the participant (Lang et al., 2020). The total exposure time was 2 minutes and 40 seconds. An eye-tracking sensor ensured continuous fixation on the images (Pernice & Nielsen, 2009). Following the visual stimulus presentation, participants completed the BRAMS for a third time (Psychometric Test 3: PT3), performed a second 3-minute dance improvisation, and concluded the protocol with a final BRAMS assessment (Psychometric Test 4: PT4) (Figure 1).

**Figure 1.**
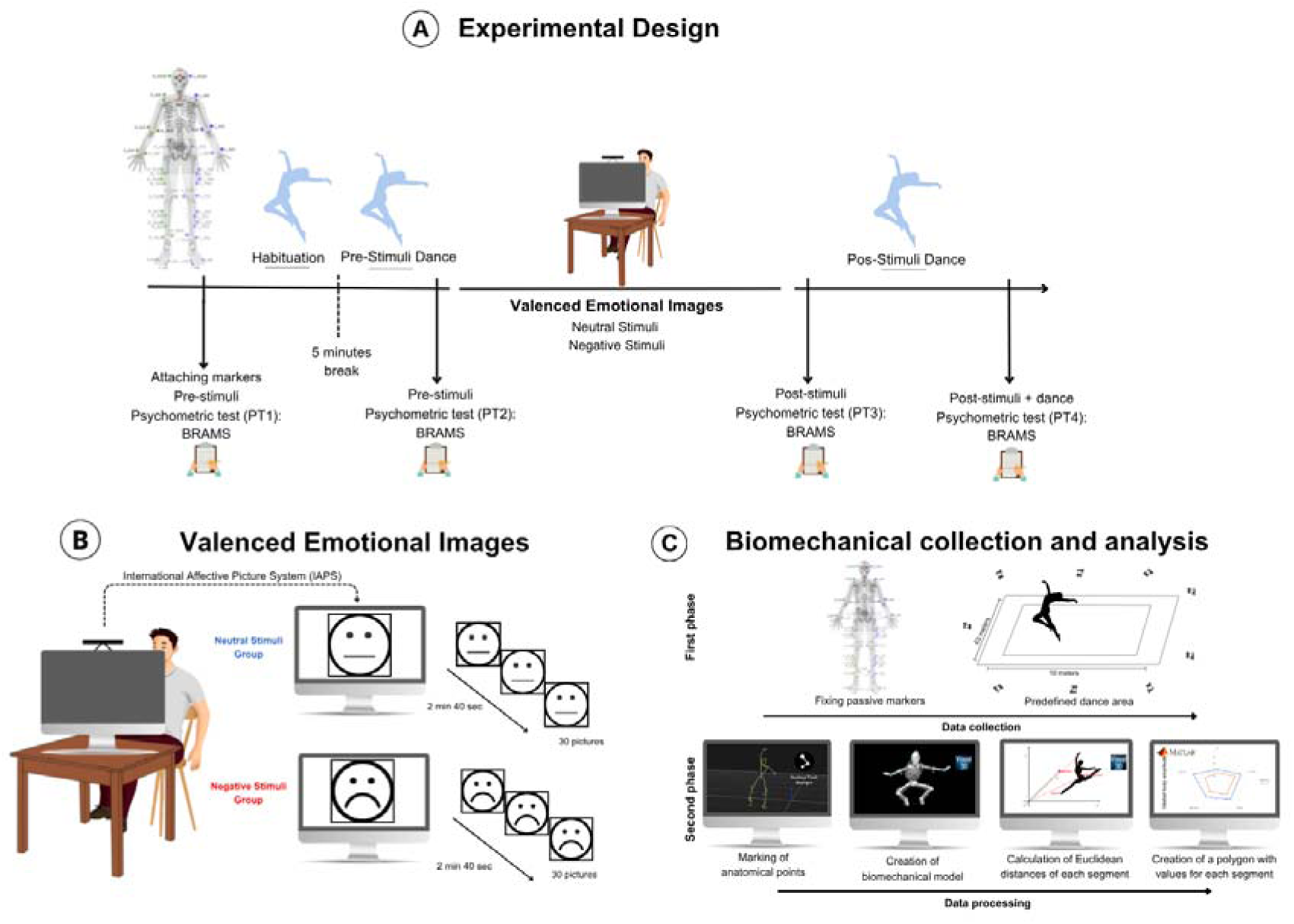
Methodological Design and Study Procedures. **(A)** Overview of the experimental protocol. Volunteers (professional dancers) were initially outfitted with passive markers and underwent Psychometric Test 1 (PT1) using the Brazilian Mood Scale (BRAMS). After familiarizing themselves with the environment, participants performed the Pre-Stimuli Dance and completed PT2. They were exposed to visual stimuli (Neutral or Negative) and subsequently completed PT3. Following the stimulus exposure, they performed Post-Stimuli Dance, and PT4 was collected. **(B)** Procedure for visual stimulus exposure. Participants were exposed to neutral (Neutral Stimuli) or negative (Negative Stimuli) visual stimuli through a 36 x 96 cm monitor placed 1 meter away. Each participant, isolated in the collection room, had their gaze tracked by an *Eyetracking* sensor to ensure attention to the monitor. The visual stimuli were from the *International Affective Picture System* (IAPS), with each image displayed for 5 seconds, for a total of 2 minutes and 40 seconds of visual exposure. All participants viewed a white screen for 5 seconds before the stimulus presentation. **(C)** Kinematic assessment of dance using the *Qualisys Track Manager* (QTM) system within the Project Automation Framework (PAF) module. Dancers performed an improvised dance in a 3 x 10-meter space for 3 minutes. Kinematic data were analysed using *Visual3D*® *software*, in which biomechanical models were created from marker positions. The kinematics of body segments was calculated using the body segment coordinate system, assuming a rigid body model. Segmental movement variables were represented as polygons, with vertices corresponding to the values of these variables for each segment (right hand, left hand, right foot, left foot, and head).

### Brazilian Mood Scale (BRAMS)

The Brazilian Mood Scale (BRAMS) is an adaptation of the Brunel Mood Scale (BRUMS), a short version of the Profile of Mood States (POMS), a measure of affective mood state fluctuation (Brandt et al., 2012; Mcnair et al., 1971; Rohlfs et al., 2008, 2023; Terry et al., 2003). BRAMS is a 24-item measure assessing the mood subscales of tension, depression, anger, vigor, fatigue, and confusion, each with 4 items. The participants indicated how they were feeling “right now” on a 5-point Likert-type scale, where 0 = not at all, 1 = a little, 2 = moderately, 3 = quite a bit, and 4 = extremely.

### Motion capture and kinematic evaluation of dance

Kinematic evaluation was performed using an optoelectronic system with 10 cameras (Qualisys) set to track movement at 300 Hz (Cappozzo et al., 2005; Robertson et al., 2004; Winter, 2009). Reflective markers were placed on the participant’s anatomical structures (Supplementary Figure), and the system reconstructed the markers’ 3D positions to create a virtual model (Supplementary Video). Calibration was performed at the beginning of each session. First, static data were collected, followed by a dynamic skiing simulation to facilitate marker recognition. Each participant danced for 3 minutes, with the first 1.5 minutes discarded from the analysis to allow for adaptation. The dance occurred without music or external interference.

### Data reduction

The collected data were transferred to the Visual3D® software, where a biomechanical model was created from the body segment coordinates. Kinematics were calculated using a rigid-body model (Santos et al., 2023), and data were filtered with a fourth-order Butterworth filter at a 6 Hz cutoff frequency (Winter, 2009). Interpolation was applied to trajectories with missing signals, with a maximum gap of 10 frames. The positions of the segmental centers of mass were estimated according to Hanavan (HANAVAN, 1964).

The distances between the extremity segments, hands, feet, and head relative to the pelvis were calculated from the difference in Euclidean distances of the segmental centers of mass. Each time series of segments-to-pelvis distances consisted of 27000 points (frames), and the range of motion of each segment was calculated as the difference between the maximum and minimum distance values of each time series. The expansion of movement for each segment was computed as the average of the segment-to-pelvis distance time series. Segmental velocity was calculated as the first derivative of the Euclidean 3D position of the segment center of mass relative to the global reference frame (i.e., laboratory). The exploration area was defined as the ellipse that encompasses 95% of the pelvis data (Duarte & Freitas, 2010).

To compute global movement variables (global movement amplitude, expansion, and velocity) as the study outcome variables, each segmental variable (segment movement amplitude, expansion, and velocity) was used to construct a five-vertex polygon for each segment (hands, feet, and head). The area of the polygon referring to the segments’ movement amplitudes corresponded to the global movement amplitude. The polygon area computation was performed to determine the global movement expansion and velocity, using the polygons corresponding to the segments’ movement expansions and velocities, respectively.

### Variables’ reliability

To assess the reliability of the dance improvisation outcome variables, the intraclass correlation coefficient (ICC) was computed for each variable measured during the familiarization and pre-stimuli dances. The results were classified according to the Fleiss scale (Fleiss, 1999). Variables with ICCs above 0.7 would be considered reliable and subjected to further statistical analyses. Only the global movement amplitude and expansion were reliable (ICCs 0,84 and 0,9, respectively). The global movement velocity was not sufficiently reliable (ICC 0,1) and was therefore excluded from the main statistical analyses.

### Statistical Analysis

The statistical analysis was conducted using linear mixed-effects models to account for both fixed and random effects. The models were fitted for both physical variables (Global Movement Amplitude and Global Movement Expansion) and psychological variables (Tension, Depression, Anger, Vigor, Fatigue, and Mental Confusion). In each model, the interaction between time and group was included as a fixed effect, and random intercepts were specified at the subject level to account for within-subject variability. The models were implemented using the lme() function from the nlme package in R. Estimated marginal means were computed for each time point and group using the emmeans package. Pairwise comparisons between groups were performed using Tukey’s post-hoc test to adjust for multiple comparisons. The p-values from these comparisons were used to assess statistical significance between groups, and the results were sorted in ascending order of p-value for clarity. To validate the models, residual diagnostics were performed. Q-Q plots were generated for each model to visually assess the normality of residuals. Additionally, the Shapiro-Wilk test was used to assess whether the residuals were normally distributed. Significance level was set at 0.05. All analyses were conducted using R software (version 4.3.2). Graphs were generated using MatLab® and GraphPad Prism® 8.2.2.

## Results

### Baseline demographic and anthropometric equivalence

Participants assigned to the Neutral versus Negative visual-stimulus conditions were comparable on all baseline demographic and anthropometric characteristics, indicating no evidence of pre-existing group differences that could confound subsequent analyses. Specifically, the groups did not differ significantly in age (Neutral: M = 31.0, SD = 12.39; Negative: M = 32.5, SD = 11.13; d = 0,13, p = 0.779), years of professional experience (Neutral: M = 12.3, SD = 13.12; Negative: M = 15.9, SD = 10.51; p = 0.507), height (Neutral: M = 165.7 cm, SD = 7.69; Negative: M = 170.1 cm, SD = 6.74; p = 0.190), or weight (Neutral: M = 63.83 kg, SD = 7.72; Negative: M = 63.34 kg, SD = 14.00; p = 0.924). Variance homogeneity assumptions were also met for each variable (all Levene’s tests p = 0.225), supporting the interpretation that the two groups were well matched at baseline (Table).

### Negative visual context selectively altered dancers’ mood states

To examine whether a negatively valenced emotional context influences the biomechanics of dance improvisation in professional dancers, we used the International Affective Picture System (IAPS) to induce either neutral or negative affective states. The IAPS is a widely used and well-validated tool for eliciting psychophysiological responses to affective stimuli (Branco et al., 2023; Lang et al., 2020; R. L. Ribeiro et al., 2004). Participants’ visual attention to the stimuli was monitored using eye-tracking, thereby helping ensure engagement with the emotional manipulation. To verify that the IAPS procedure altered participants’ mood, particularly in the negative-stimulus condition, we administered the psychometric test (PT) Brazilian Mood Scale (BRAMS), a Brazilian adaptation of the Brunel Mood Scale (BRUMS), itself a brief version of the Profile of Mood States (POMS) (Bevilacqua et al., 2019; Searight & Montone, 2017; Shacham, 1983; Terry et al., 1999). Initially, professional dancers were randomly assigned to either the Neutral Stimuli or Negative Stimuli group, and mood was first assessed (PT1) before the experimental procedures (Figure 1). At PT1, dancers in the Negative Stimuli group showed higher scores for tension (d= 3,94, p = 0.005) (Figure 2A) and depression (d = 2,99, p = 0.03) (Figure 2B) relative to the Neutral Stimuli group. There were no differences between groups for anger (Figure 2C), vigor (Figure 2D), fatigue (Figure 2E), and mental confusion (Figure 2F).

**Figure 2.**
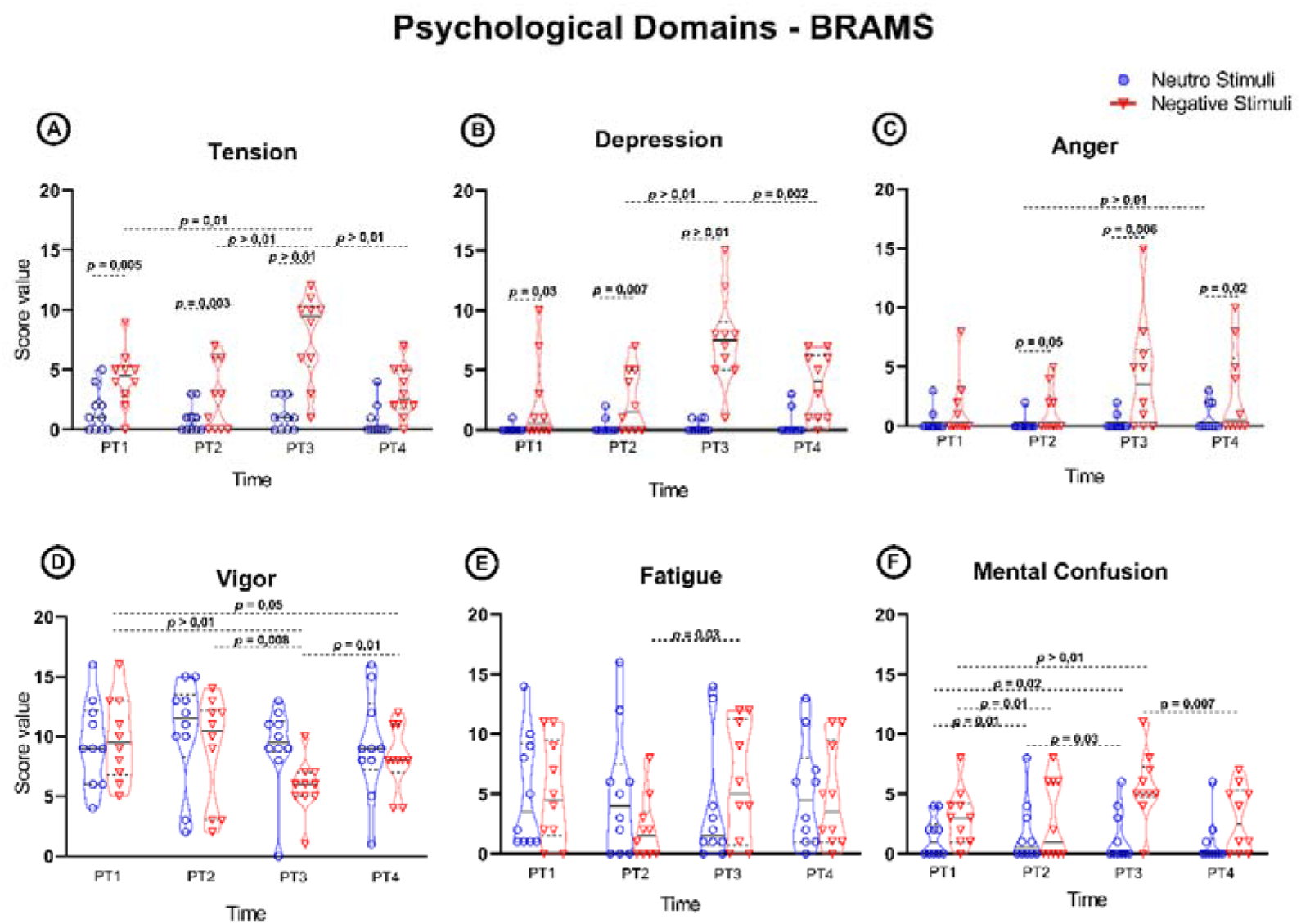
Mood responses to neutral and negative visual stimulation and subsequent dance. Within-group analyses showed that, in the Negative Stimuli condition, exposure to negatively valenced images (PT2 vs. PT3) increased tension, depression, and fatigue following negative-valence visual stimulation. Conversely, the Negative Stimuli group exhibited decreased vigor in response to negative-valence visual stimuli (PT2 vs. PT3). The Neutral Stimuli group showed no changes across most domains, except for a decrease in mental confusion (PT2 vs. PT3) following neutral-valence visual stimulation. After the Post-Stimuli dance, there was a reduction in tension, depression, and mental confusion, and an increase in vigor (PT3 vs. PT4). Between-group comparisons indicated baseline differences at PT1, with differences between the Neutral Stimuli dancers and the Negative Stimuli dancers in terms of tension, depression, anger, and mental confusion. At PT2, there were differences between the Neutral Stimuli dancers and the Negative Stimuli dancers in terms of tension, depression, and anger. There were significant differences in tension, depression, and anger at PT3. At PT4, a difference in anger was observed between Neutral Stimuli dancers and Negative Stimuli dancers. Blue = Neutral Stimuli dancers; Red = Negative Stimuli dancers; PT = Psychometric Test 1, 2, 3, and 4 = mood assessment time points. Data were analyzed using the Generalized Estimating Equations method, with experimental groups and dances as fixed effects and individual dancers as random effects. Post hoc pairwise comparisons were conducted among all experimental groups. Statistical significance was determined as p < 0.05.

Following room habituation and completion of the first dance improvisation (Pre-Stimuli dance), participants completed the second mood assessment (PT2). Overall, Pre-Stimuli dance did not affect the dancers’ mood. At PT2, dancers in the Negative Stimuli group still reported higher scores for tension (d = 4,27, p = 0,003) (Figure 2A), depression (d = 3,78, p = 0,007) (Figure 2B), and anger (d = 2,8, p = 0,05) (Figure 2C) than those in the Neutral Stimuli group. As at PT1, no between-group differences were observed for vigor (d = 0,28, p = 0,84) (Figure 2D) and fatigue (d = 0,52, p = 0,71) (Figure 2E). Within-group comparisons between PT1 and PT2 further indicated relative mood stability across most psychometric domains. The only significant change was an increase in mental confusion from PT1 to PT2 in both the Neutral Stimuli (d = 2,13, p = 0,01) and the Negative Stimuli group (d = 2,51, p = 0,01) (Figure 2F). No significant PT1-PT2 changes in tension, depression, anger, vigor, or fatigue were observed in either group (Figure A-E).

We next examined whether the valenced visual context altered dancers’ mood states. After image exposure, participants in both the Neutral Stimuli and Negative Stimuli conditions completed the third mood assessment (PT3) (Figure 1). In the Neutral Stimuli condition, there were no significant changes in tension, depression, anger, vigor, or fatigue from PT2 to PT3 (Figure 2A-E). However, we observed a reduction in mental confusion (d = 0,89, p = 0,03) (Figure 2F). In contrast, dancers exposed to negatively valenced images showed a change in mood from PT2 to PT3, characterized by increases in tension (d = 3,44, p < 0,001) (Figure 2A), depression (d = 1,97, p < 0,001) (Figure 2B), and fatigue (d = 2,86, p = 0,03) (Figure 2E), alongside a reduction in vigor (d = 2,89, p = 0,008) (Figure 2D). No significant PT2–PT3 changes were observed for anger (Figure 2C) or mental confusion (Figure 2F). At PT3, the Negative Stimuli group also reported higher tension (d = 6,01, p < 0,001) (Figure 2A), depression (d = 7,9, p < 0,001) (Figure 2B), anger (d= 3,91, p = 0,006) (Figure 2C), vigor (d = 4,87, p < 0,001) (Figure 2D), and mental confusion (d = 4,04, p = 0,004) (Figure 2F) than the Neutral Stimuli group. No between-group differences were detected at PT3 for fatigue (p = 0,28) (Figure 2E).

Following the second dance improvisation (Post-Stimuli dance), participants completed the fourth mood assessment (PT4) (Figure 1). In the Neutral Stimuli group, no significant changes were observed from PT3 to PT4 across the assessed mood domains (Figure 2). In contrast, dancers in the Negative Stimuli group showed significant reductions in tension (d = 5,11, p < 0,001) (Figure 2A), depression (d = 3,47, p = 0,002) (Figure 2B), vigor (d = 3,22, p = 0,01) (Figure 2D), and mental confusion (d = 4,04, p = 0,007) (Figure 2F) from PT3 to PT4, whereas anger (d = 0,5, p = 0,5) (Figure 2C) and fatigue (d = 1,24, p = 0,06) (Figure 2E) did not change significantly. At PT4, the only remaining between-group difference was anger, which remained higher in the Negative Stimuli group than in the Neutral Stimuli group (d = 3,2, p =0,02) (Figure 2C).

### Negative visual context selectively altered the biomechanics of improvised dance

Using the IAPS manipulation in combination with repeated BRAMS assessments, we first showed that negatively valenced visual stimuli altered dancers’ mood states (Figure 2). We then tested whether this affective manipulation was accompanied by changes in the biomechanics of improvised dance. In the Neutral Stimuli group, no significant differences were observed between Pre-Stimuli dance and Post-Stimuli dance for either global movement amplitude (Figure 3A) or the global movement expansion (Figure 3B). On the other hand, dancers in the Negative Stimuli group showed a significant reduction in the global movement amplitude during Post-Stimuli dance relative to Pre-Stimuli dance (d = 1,28, p = 0,001) (Figure 3A), while the global movement expansion remained unchanged (Figure 3B). No between-group differences were detected for these variables in either Pre-Stimuli dance or Post-Stimuli dance (Figure 3).

**Figure 3.**
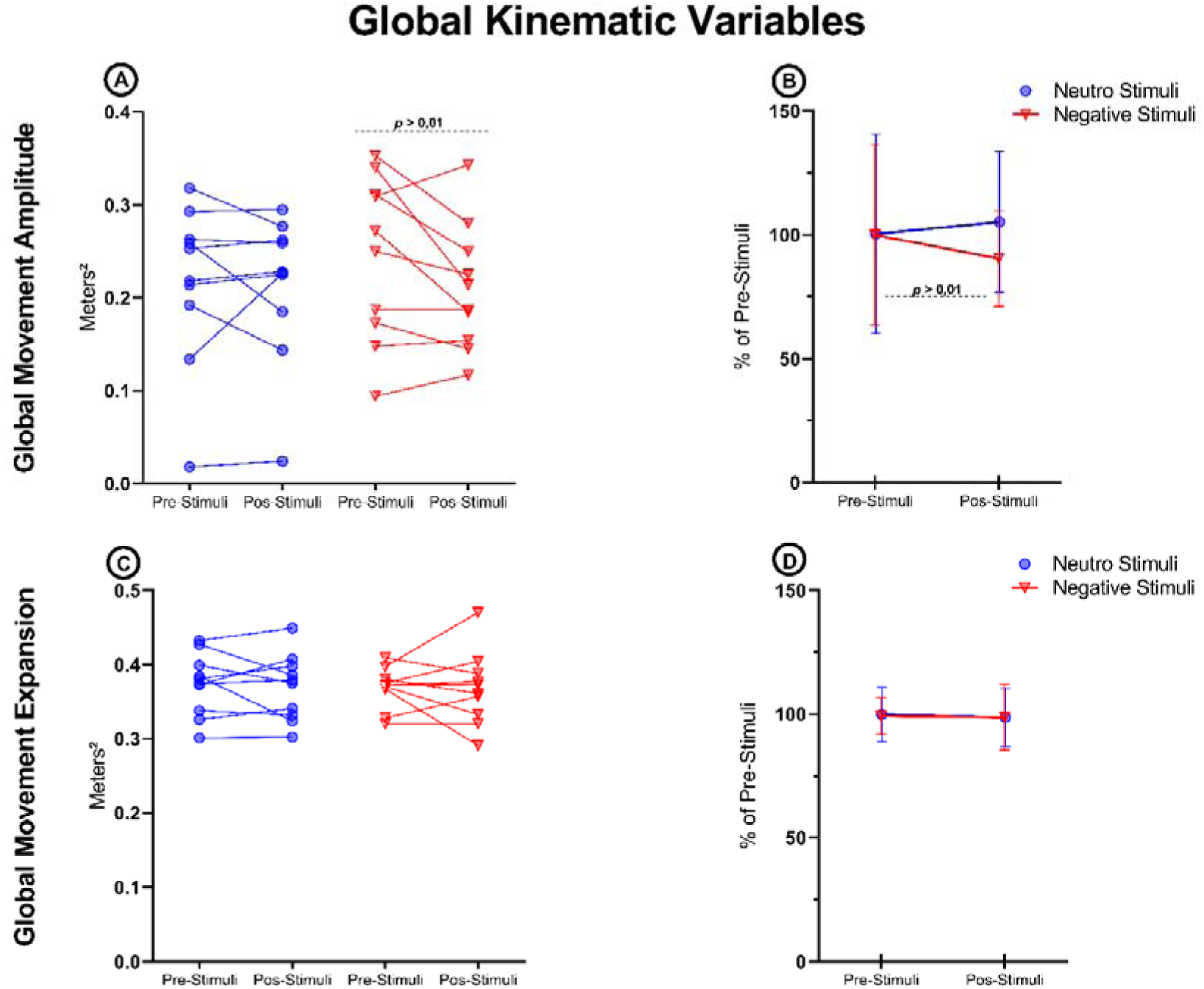
Global kinematic variables during dance improvisation before and after emotional valence visual stimuli. (A) Global average amplitude: A significant reduction in amplitude was observed only in the Negative Stimuli group following the negative visual stimulus. (B) Global average expansion: No significant differences were observed between groups (Neutral Stimuli vs. Negative Stimuli) or within groups over time (Pre-Stimulus Dance vs. Post-Stimulus Dance). Data were analyzed using the Generalized Estimating Equations method, with experimental groups and dances as fixed effects and individual dancers as random effects. Post hoc pairwise comparisons were conducted among all experimental groups. Statistical significance was determined as p < 0.05.

## Discussion

In this study, we tested the hypothesis that dance movements not only express what the dancer intends to communicate but also carry an additional signature of the dancer’s current affective state. Here we found that exposure to negatively valenced images altered both psychological state and subsequent movement biomechanics, indicating that transient emotional context can measurably shape improvised dance behavior. Negative-valence IAPS exposure reliably altered mood and was followed by a measurable reduction in global movement amplitude during improvised dance, whereas global movement expansion remained unchanged. Together, these findings indicate that transient negative affect can influence the biomechanics of improvised dance, even in highly trained performers.

A key manipulation check was the PT2–PT3 change in mood domains, which occurred specifically in the Negative Stimuli group. Following exposure to negatively valenced images, these dancers reported higher tension, depression, and fatigue, alongside reduced vigor, while the Neutral Stimuli group showed no comparable shifts across these domains and only a decrease in mental confusion. Moreover, at PT3, the Negative Stimuli group scored higher than the Neutral group on tension, depression, and anger, indicating a condition-specific shift toward a more negatively valenced mood profile. Taken together, these patterns suggest that the negative visual context produced the intended shift toward a more negatively valenced mood state. The ability to reliably induce such affective changes is a defining strength of standardized emotional picture paradigms, which provide stimuli with normative affective properties and support reproducible emotion induction across samples and cultural contexts (Branco et al., 2023; Lang et al., 2020; R. L. Ribeiro et al., 2004).

We further observed that following negative IAPS exposure, dancers in the Negative Stimuli group showed a reduction in global movement amplitude from Pre-Stimuli dance to Post-Stimuli dance, whereas global movement expansion remained unchanged. No comparable changes were observed in the Neutral Stimuli group for either kinematic indicator. Together, these findings suggest that negatively valenced visual context selectively reduced the magnitude of movement of the body’s extremities during free improvisation, rather than positioning the extremities closer or farther from the body center. This pattern is consistent with a broader literature demonstrating that affective visual cues can influence motor control, including postural stability and movement initiation in static and cyclic tasks (Benesova et al., 2026; Bouman et al., 2015; Brandão et al., 2016; Coombes et al., 2005; D’Attilio et al., 2013; Stins & Beek, 2007), and can also modulate motor-system readiness and corticospinal excitability during picture viewing (Coelho et al., 2010; Hajcak et al., 2007; van Loon et al., 2010). In an ecological performance setting, a reduction in movement amplitude may reflect an adaptive motor strategy in response to negative affect, potentially involving increased muscular co-contraction, altered action readiness, or a more protective movement style (Adkin & Carpenter, 2018; Doumas et al., 2018). Importantly, our results show that transient emotional context can shape movement expression in a complex, whole-body improvisational behavior relevant to artistic performance. The lack of change in global expansion further suggests that negative mood induction altered movement vigor without substantially shifting the dance’s overall spatial “footprint.” One plausible interpretation is that professional dancers maintained broad spatial organization while reducing energetic scaling, consistent with selective effects of affect on movement intensity rather than on movement topology. This dissociation is noteworthy because it implies that the immediate affective context may be expressed primarily through the body’s movement (intensity and amplitude) rather than where it moves (overall spatial deployment), potentially preserving externally constrained or socially learned performance norms despite internal emotional perturbation. Movement velocity could help test this hypothesis, particularly regarding movement intensity; however, this variable was not sufficiently reliable.

An additional pattern was that, in the absence of negative stimulation, psychological domains remained broadly stable across PT1 to PT4, suggesting that dance improvisation alone did not change mood in a low-stress setting. However, after negative stimulation, dancers showed partial normalization from PT3 to PT4 (reductions in tension, depression, and confusion, with increased vigor), suggesting that dance may contribute to affective recovery following emotional perturbation. This aligns with embodied and action-based accounts, in which modifying movement patterns can contribute to emotion regulation (Aguiar et al., 2016; Anderson et al., 2014). Conceptually, this supports the idea that dance may not only reflect affect, but also reshape it, especially following acute affective perturbations. However, because we did not include the “negative stimulation without subsequent dance” condition, we cannot disentangle spontaneous recovery from dance-mediated regulation.

More broadly, these findings speak to central questions on how immediate affective cues in the surrounding environment shape emotion, creativity, and skilled action. Across artistic domains, mood and affect have been linked to creative performance and artistic choices (Baas et al., 2008; He, 2023; McPherson et al., 2014; To et al., 2012; Verhaeghen et al., 2005), including emotionally targeted improvisation paradigms showing that affective intent and context can shape expressive structure and underlying neurocognitive dynamics (Gomez & Danuser, 2007; McPherson et al., 2016). In parallel, sport and performance research demonstrates that anxiety and affective load can disrupt precision, alter attentional control, and impair outcomes under pressure (Horikawa & Yagi, 2012; Nibbeling et al., 2012; Wilson, 2008; Yu, 2015). In addition, research using emotional pictures shows that valence/arousal modulate cognitive-motor output and kinematics (Benesova et al., 2026; Coombes et al., 2005; Esteves et al., 2016; Larson et al., 2006; Stins & Beek, 2007). Our data extend these principles to improvised dance. Negative emotional context, induced by standardized visual stimuli, shifted the internal state and selectively reduced movement amplitude in the area, consistent with the idea that what audiences perceive as “expression” may partly reflect the dancer’s state rather than only the dancer’s intention trained during the work process. Or maybe it reflects a change in the dancer’s intention.

Some challenges and limitations must be considered. First, one challenge in our results is baseline imbalance across psychological domains. The Negatively Stimulated group already differed from the Neutral Stimulated group at arrival (PT1) and remained different at PT2 in domains including tension and depression (and anger at PT2) prior to the emotional-stimulus manipulation. This pre-existing elevation in negative mood could have reduced sensitivity to detect between-group biomechanical differences, even though within-group changes following negative stimulation were evident. Future studies could mitigate this by using stratified randomization based on baseline mood, including baseline mood as a covariate, and/or larger sample sizes. Second, our sample comprised professional dancers, and training may shape affective processing and the perception of expressiveness, potentially moderating the magnitude or form of emotion-movement coupling (Di Mauro et al., 2018; Park et al., 2014). Furthermore, the sample size limited our ability to test sex-specific effects, which may be relevant given reported sex differences in electrophysiological responses to IAPS content (Kemp et al., 2004). Third, some movement features (e.g., speed, exploration area) were excluded due to reliability constraints, narrowing inference to global amplitude and expansion area metrics. Fourth, the study design did not include a post-stimulus “negative stimulation without subsequent dance” condition, which prevented us from distinguishing spontaneous recovery from dance-facilitated regulation following negative stimulation. Finally, while the laboratory environment enabled control of the visual manipulation, supported by eye-tracking verification of attention, it also simplified the ecological context of dance by excluding music. Real-world dance is inherently multisensory, and auditory input can both structure movement and modulate emotion. Future research should test how multisensory integration shapes biomechanical responses to affective contexts, including whether music buffers against reductions in movement amplitude under negative visual stimulation.

The present study underscores a practical and conceptual point: understanding how emotion is involved in artistic movement is essential not only for theories of creativity and communication but also for artists’ well-being. If dancers’ biomechanics are shaped by their emotional state, even when they are not explicitly “performing” that emotion, then psychological stressors in professional contexts may influence intention, expressive quality, and physical load. Clarifying these links can inform training environments, performance preparation, and mental-health support strategies that help artists sustain both creativity and health over time.

## Declarations

## Acknowledgements

To the young scientists at the NeuroDEv laboratory, thank you for your daily support. To our colleagues, we appreciate our scientific discussions and the assistance you provided in times of urgency. And a special acknowledgment to the often “invisible” yet indispensable members of our university community - the technicians, cleaning personnel, security team, restaurant staff, and administration professionals - your contributions are the bedrock of our research endeavor.

## Ethics approval

This study was performed in line with the principles of the Declaration of Helsinki. This study was approved by the Research Ethics Committee of the Federal University of Minas Gerais under protocol number 5.697.800 (CAAE - 58101121.3.0000.5149).

## Informed consent

All volunteers signed an Informed Consent Form (ICF), acknowledging the procedures they would undergo and agreeing to participate in the study.

## Availability of data and materials

Data or materials for the experiments are available upon request

## Authors’ contributions

BCBM: Conceptualization, Methodology, Investigation, Data Curation, Writing - Original Draft, Visualization

TRS: Conceptualization, Methodology, Validation, Resources, Writing - Review & Editing, Supervision, Project administration, Funding acquisition

GSO: Investigation

JBPN: Investigation

CESS: Investigation

CBP: Investigation

JBLS: Validation, Writing - Review & Editing, Supervision

LAOAN: Methodology, Validation

PAA: Methodology, Validation

RPS: Formal analysis

BRS: Conceptualization, Methodology, Validation, Resources, Writing - Review & Editing, Visualization, Supervision, Project administration, Funding acquisition

---

**Table.**
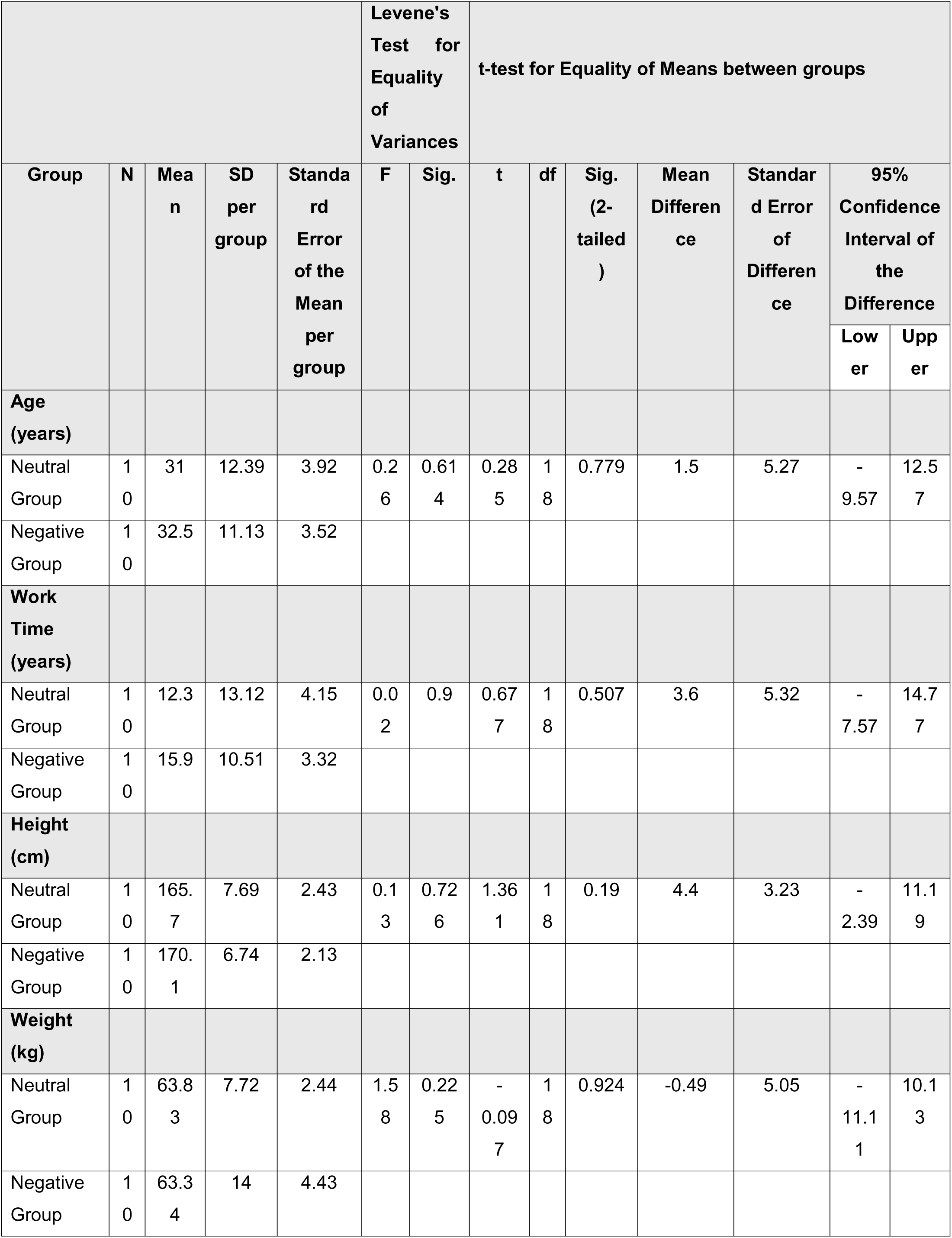
Anthropometric characteristics and professional profile of the dancers. Data are expressed as Mean ± Standard Deviation (SD). Levene’s Test was used to verify the equality of variances, and Student’s independent-samples t-test was applied to compare the means between the Control Group (n=10) and the Intervention Group (n=10). No statistically significant differences were observed (p > 0.05) regarding age, professional work time, height, or weight, confirming the initial homogeneity between the groups.

