## Supplementary Figure for "Negative emotional visual stimuli alter specific improvised dance biomechanics in professional dancers"

### Qualisys PAF package: Functional Assessment marker set - Upper body

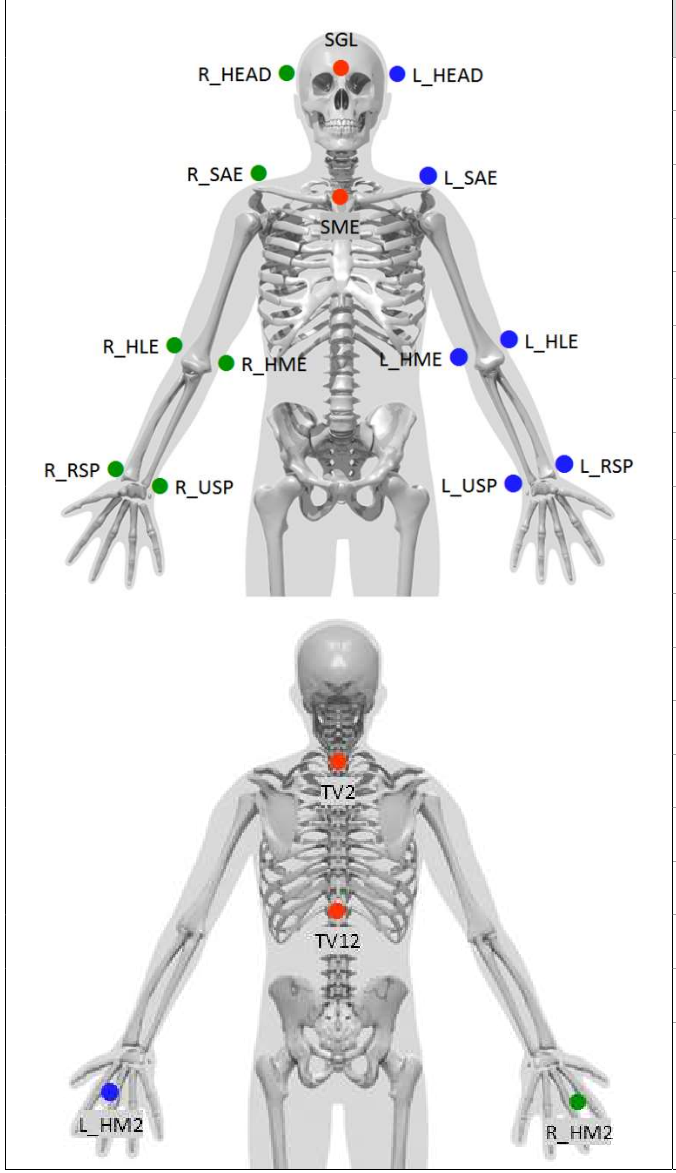


| Name | Ref. [1] | Location | Static (18) | Dyn. (18) |
|---|---|---|---|---|
| L_HEAD | | On headband, just above ear | X | X |
| R_HEAD | | On headband, just above ear | X | X |
| SGL | SGL | On headband, Forehead | X | X |
| SME | SME | Sternum | X | X |
| TV2 | TV2 | Spine, 2nd Thoracic Vertebra | X | X |
| TV12 | TV12 | Spine, 12th Thoracic Vertebra | X | X |
| L_SAE | SAE | Shoulder | X | X |
| L_HLE | HLE | Elbow (outside) | X | X |
| L_HME | HME | Elbow (inside) | X | X |
| L_RSP | RSP | Wrist (thumb side) | X | X |
| L_USP | USP | Wrist (pinkie side) | X | X |
| L_HM2 | HM2 | Hand (basis of Forefinger) | X | X |
| R_SAE | SAE | Shoulder | X | X |
| R_HLE | HLE | Elbow (outside) | X | X |
| R_HME | HME | Elbow (inside) | X | X |
| R_RSP | RSP | Wrist (thumb side) | X | X |
| R_USP | USP | Wrist (pinkie side) | X | X |
| R_HM2 | HM2 | Hand (basis of Forefinger) | X | X |

[1] Sint Jan, S. Van (2007). Color Atlas of Skeletal Landmark Definitions. Guidelines for Reproducible Manual and Virtual Palpations. Edinburgh: Churchill Livingstone.

Copyright © Qualisys AB

#### Qualisys PAF package: Functional Assessment marker set - Lower body

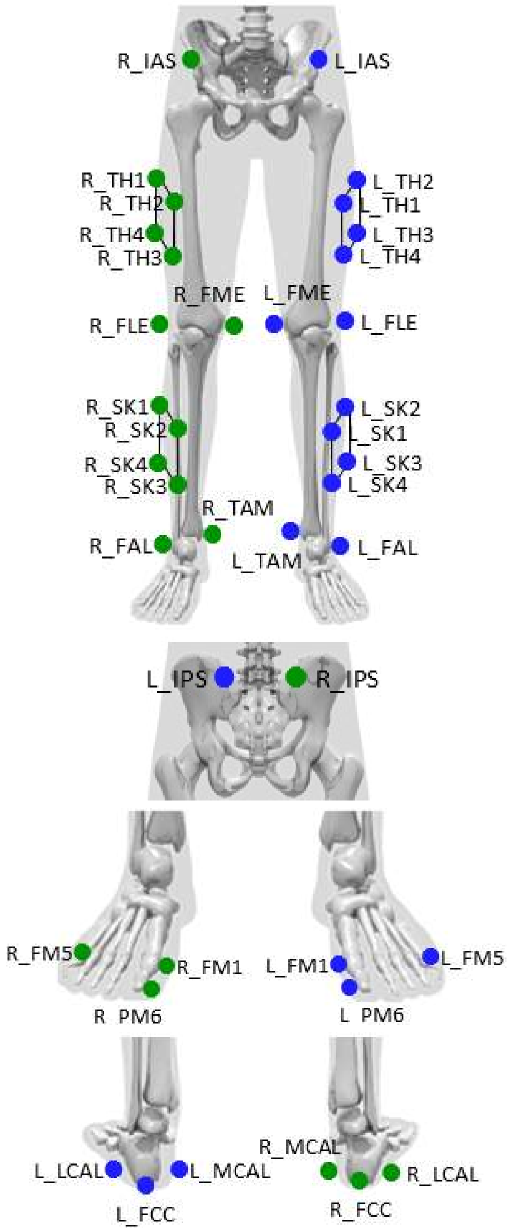


| Name | Ref. [1] | Location | Static (28) | Dyn. (28) |
|---|---|---|---|---|
| L_IAS | IAS | Anterior superior iliac spine | X | X |
| L_IPS | IPS | Posterior superior iliac spine | X | X |
| R_IPS | IPS | Posterior superior iliac spine | X | X |
| R_IAS | IAS | Right anterior superior iliac spine | X | X |
| L_TH1-4 | | Cluster | X | X |
| L_FLE | FLE | Lateral epicondyle | X | X |
| L_FME | FME | Medial epicondyle | X | X |
| L_SK1-4 | | Cluster | X | X |
| L_FAL | FAL | Lateral prominence of the lateral malleolus | X | X |
| L_TAM | TAM | Medial prominence of the medial malleolus | X | X |
| L_FCC | FCC | Aspect of the Achilles tendon insertion on the calcaneus | X | X |
| L_LCAL | | Lateral calcaneus | X | X |
| L_FM5 | FM5 | Dorsal margin of the fifth metatarsal head | X | X |
| L_PM6 | PM6 | Proximal medial phalanx of the big toe | X | X |
| L_FM1 | FM1 | Dorsal margin of the first metatarsal head | X | X |
| L_MCAL | | Medial calcaneus | X | X |
| R_TH1-4 | | Cluster | X | X |
| R_FLE | FLE | Lateral epicondyle | X | X |
| R_FME | FME | Medial epicondyle | X | X |
| R_SK1-4 | | Cluster | X | X |
| R_FAL | FAL | Lateral prominence of the lateral malleolus | X | X |
| R_TAM | TAM | Medial prominence of the medial malleolus | X | X |
| R_FCC | FCC | Aspect of the Achilles tendon insertion on the calcaneus | X | X |
| R_LCAL | | Lateral calcaneus | X | X |
| R_FM5 | FM5 | Dorsal margin of the fifth metatarsal head | X | X |
| R_PM6 | PM6 | Dorsal aspect of the second metatarsal head | X | X |
| R_FM1 | FM1 | Dorsal margin of the first metatarsal head | X | X |
| R_MCAL | | Medial calcaneus | X | X |

[1] Sint Jan, S. Van (2007). Color Atlas of Skeletal Landmark Definitions. Guidelines for Reproducible Manual and Virtual Palpations. Edinburgh: Churchill Livingstone.

Copyright © Qualisys AB
